# The curse of the red pearl: a fibroblast specific pearl-necklace mitochondrial phenotype caused by phototoxicity

**DOI:** 10.1101/2024.09.05.611420

**Authors:** Irene MGM Hemel, Kèvin Knoops, Carmen López-Iglesias, Mike Gerards

**Affiliations:** Maastricht Centre for Systems Biology (MaCSBio), Maastricht University, Maastricht, 6229 EN, the Netherlands; Microscopy CORE lab, Maastricht University, Maastricht, 6229 ER, the Netherlands

## Abstract

The dynamic nature of mitochondria makes live cell imaging an important tool in mitochondrial research. Although imaging using fluorescent probes is the golden standard in studying mitochondrial morphology, these probes might introduce a-specific features. In this study, live cell fluorescent imaging was applied to investigate a pearl-necklace shaped mitochondrial phenotype that arises when mitochondrial fission is restricted. In this fibroblast specific pearl-necklace phenotype, constricted and expanded mitochondrial regions alternate. Imaging studies revealed that the formation time of this pearl-necklace phenotype differs between laser scanning confocal, widefield and spinning disk confocal microscopy. We found that the phenotype formation correlates with the excitation of the fluorescent probe and is the result of phototoxicity. Interestingly, the phenotype only arises in cells stained with red mitochondrial dyes. Serial section electron tomography pearl-necklace mitochondria revealed that the mitochondrial membranes remained intact, while the cristae structure was altered. Furthermore, filaments and ER were present at the constricted sites. This study illustrates the importance of considering experimental conditions for live cell imaging to prevent imaging artefacts that can have a major impact on the obtained results.

## 1. Introduction

Mitochondria are cellular organelles crucial for intracellular energy production, calcium homeostasis, apoptosis and various other processes [1]. In order to meet these variable demands, they adapt to cellular requirements, which is possible through the different mitochondrial dynamics processes. Two of these processes are fission and fusion. Mitochondrial fission is important for mitochondrial turnover and transport. The division occurs following pre-constriction at ER-mitochondrial contact sites and depends on the recruitment of Drp1 by various adaptor proteins, which form a ring structure around the mitochondria and result in the separation of the membranes [2-4]. Mitochondrial fusion allows exchange of mitochondrial content and is the result of subsequent outer and inner membrane fusion, executed by Mfn1/2 and OPA1 respectively [5, 6]. The balance between these processes is crucial to maintain a healthy mitochondrial pool.

Research into mitochondrial dynamics relies on imaging techniques to visualize mitochondrial morphology, which can provide insight into the structural alterations occurring from changes in mitochondrial dynamics processes. Fluorescent probes are frequently used to visualize mitochondrial morphology, for instance through dyes like MitoTracker [7]. A commonly applied approach to study mitochondrial dynamics is the direct assessment of mitochondrial morphology changes, often done through quantification of changes in mitochondrial shape between the various conditions [6, 8-10]. Alternatively, photoactivable or photoconvertable fluorescent proteins can be used to track the diffusion of signal across mitochondria as a measure of fusion and fission events [11-13]. Several factors are important for reliable assessment of the mitochondrial morphology, including the application of unbiased imaging and utilisation of an accurate quantification method [14]. Unlike imaging of fixed cells, live imaging allows the observation of changes in real time and the visualisation/tracking of mitochondrial motility, fission and fusion events [15].

While live cell imaging is the preferred approach to study mitochondria, there are still significant challenges. For instance, fluorescent live cell imaging could induce ROS production in cells, which is a major cause of phototoxicity [16]. Mitochondrial dynamics is specifically affected by phototoxicity [17, 18] and this should be taken into consideration when analysing mitochondrial morphology with live cell imaging. In this study, we describe a cell type specific pearl-necklace shaped mitochondrial phenotype, induced in response to laser excitation of red mitochondrial dyes during live cell imaging. Our data contributes new information highlighting the importance of considering experimental conditions for live cell imaging to prevent imaging artefacts that can have a major impact on the obtained results. Furthermore, we applied serial electron tomography to show the effect of phototoxicity in high resolution and show the benefit of this approach in understanding structural changes in mitochondria.

## 2. Results

### 2.1 Fission knockdown results in a pearl-necklace like phenotype in fibroblasts

Depletion of Drp1 in cells is known to induce mitochondrial elongation, due to an inhibition of fission [19]. However, we observed an aberrant mitochondrial phenotype upon Drp1 knockdown in normal human dermal fibroblasts (nHDFs), which was not present in cells with a non-targeting (GFP) knockdown (Figure 1A/B). While the mitochondria were elongated, as expected, they also formed a pearl-necklace (PN) like phenotype with alternating constrictions and expansions (Figure 1B). This phenotype was mostly present in the periphery of cells, however in more severe cases it was also observed around the nucleus. To confirm the observed PN-phenotype was not a cell line specific abnormality, Drp1 knockdowns were created in three other dermal fibroblast cell lines (c0388, c0407, c2244), which all presented with the same phenotype (Figure S1). Contrastingly, Drp1 knockdown in other cell types resulted in mitochondrial elongation without PN-phenotype formation (Figure 1C-F), pointing towards a cell type specific phenomenon.

**Figure 1:**
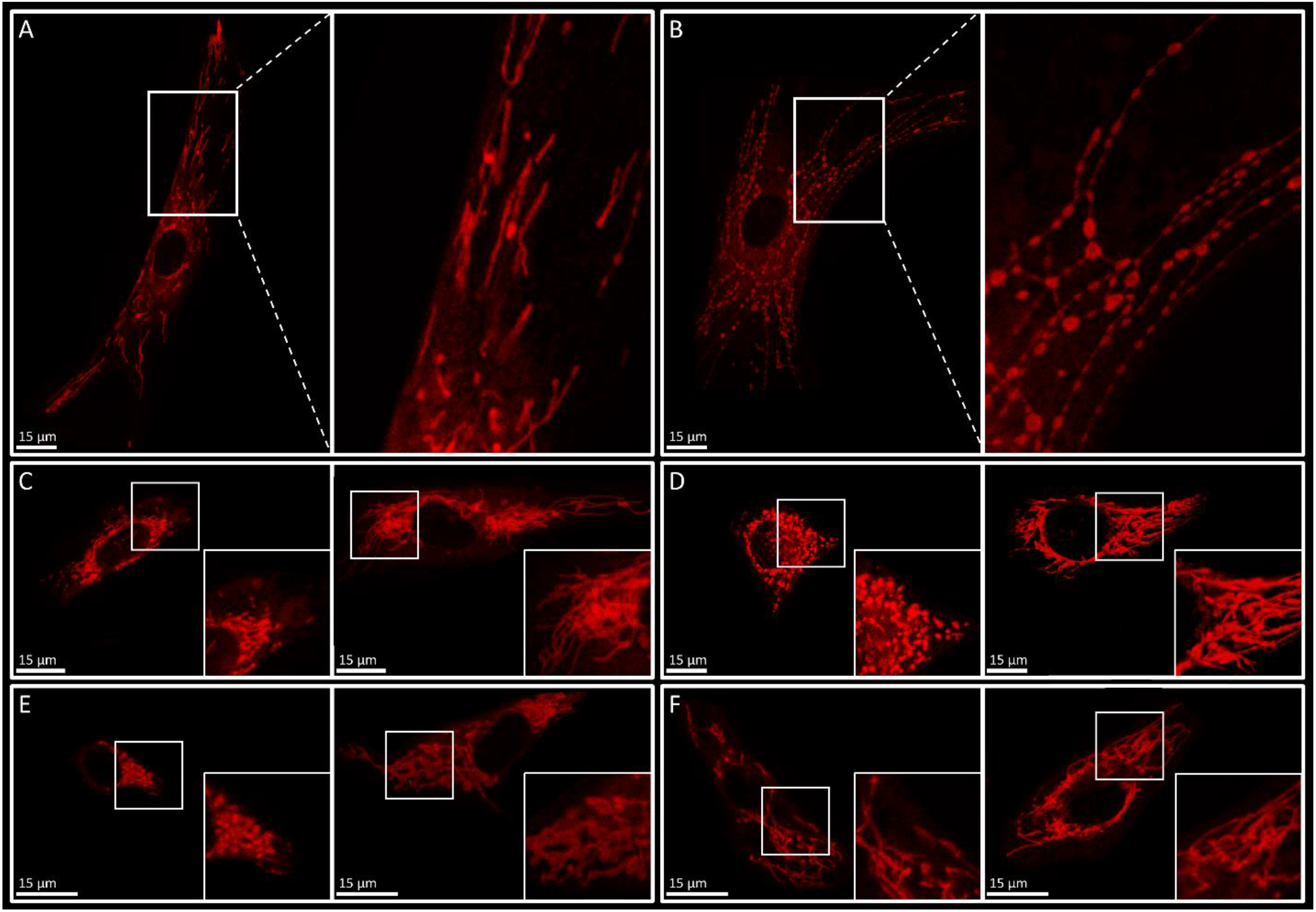
Mitochondrial morphology in a cell with a non-targeting (GFP, left) or Drp1 (right) knockdown. A/B) In addition to mitochondrial elongation normal human dermal fibroblasts present with a pearl-necklace like phenotype upon Drp1 knockdown, which is not present in control cells (enlarged squares). C-F) Other examined cell types show mitochondrial hyperfusion, but no pearl-necklace like phenotype following Drp1 knockdown. C) HUVEC cells. D) HeLa cells. E) HepG2 cells. F) Mesoangioblasts.

Fibroblasts with knockdown of one of four other proteins involved in fission (MFF, MIEF1, MIEF2 or FIS1) also presented with the PN-phenotype, indicating that the formation of this phenotype is the result of a decrease in fission in general, not just due to Drp1 knockdown. However, the amount of cells presenting with the PN-phenotype and severity of the phenotype differed between the various knockdowns. Depletion of Drp1 resulted in PN-phenotype in over 90 percent of cells, while MFF knockdown only presented the phenotype in around 35 percent of cells (Figure 2A). To confirm that the lower levels of PN-phenotype upon MFF knockdown arose from the change in gene and not a change in efficiency, the knockdown efficiency was assessed for Drp1 and MFF. Drp1 knockdown resulted in a 70% decrease in gene expression, while MFF knockdown resulted in a 90% decrease (Figure S2). This shows that the mechanism behind the formation of the PN-phenotype is not equally dependent on all fission related factors. Furthermore, to assess whether the phenotype was only present upon fission depletion, or also following increased fusion, fibroblasts were exposed to starvation, which is known to increase the mitochondrial fusion activity. Mild starvation resulted in mitochondrial elongation, without PN-phenotype, while PN-phenotype was present in fibroblasts exposed to extreme starvation (Figure 2B/C). Combined this data indicates the PN-phenotype is the result of an imbalance in mitochondrial fission and fusion.

**Figure 2:**
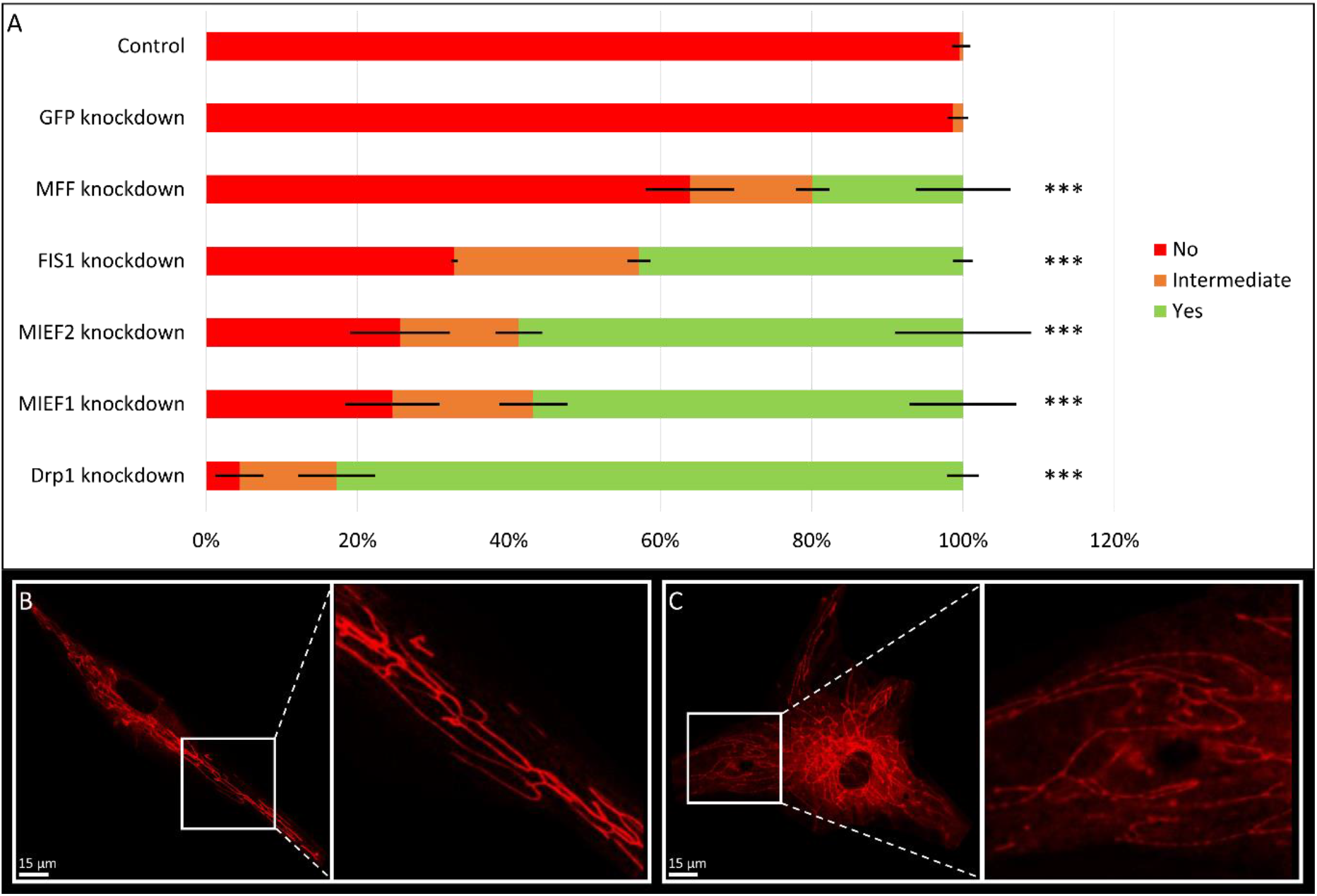
Formation of the pearl-necklace (PN) phenotype in different conditions. A) Knockdown of six fission factors all result in the formation of the PN-phenotype, with different severities, while a non-targeting (GFP) knockdown does not. Drp1 knockdown results in the most severe phenotype, with pearl structures in over 90% of cells, while MFF knockdown only results in the PN-phenotype in ∼35% of cells. ***p < 0.001 (Independent sample T-test, significance level adjusted for multiple testing). B) Following mild starvation no pearls are formed, while mitochondrial morphology is elongated. C) Upon extreme starvation, mitochondria elongate and form the PN-phenotype.

### 2.2 Formation of the pearl-necklace phenotype is the result of fluorescent laser exposure

To confirm the presence of the PN-phenotype on a different microscope system, a CorrSight, with both a widefield and a spinning-disc confocal imaging mode, was used to visualize fibroblasts with a Drp1 knockdown. Strikingly, the observed phenotype differed significantly depending on the type of microscope used. Application of laser scanning confocal microscopy (LSCM) caused rapid formation of the phenotype, resulting in visible pearls within seconds of the start of exposure, which stayed similar over the course of imaging (Figure 3A). Contrastingly, with widefield fluorescent microscopy and spinning disk confocal microscopy (SDCM) pearls were only formed around 20 seconds after the onset of exposure, while the phenotype became more severe throughout the first minute of exposure (Figure 3A). Since one of the main differences between these microscopes is the amount of light exposure, we hypothesized that the formation of pearls is caused by the excitation of the fluorescent dye with a laser. To confirm this, we imaged live cells using LSCM, followed by fixation and subsequent imaging of cells in the same well. While laser exposure in live cells resulted in rapid pearl formation, the fixed cells did not show the PN-phenotype (Figure 3B). Additionally, upon discontinuation of the laser exposure in live cells the mitochondria returned to a normal elongated shape (Figure S3). Combined, this shows that the PN-phenotype is a dynamic reversible adaptation of mitochondria as the result of continued laser exposure.

**Figure 3:**
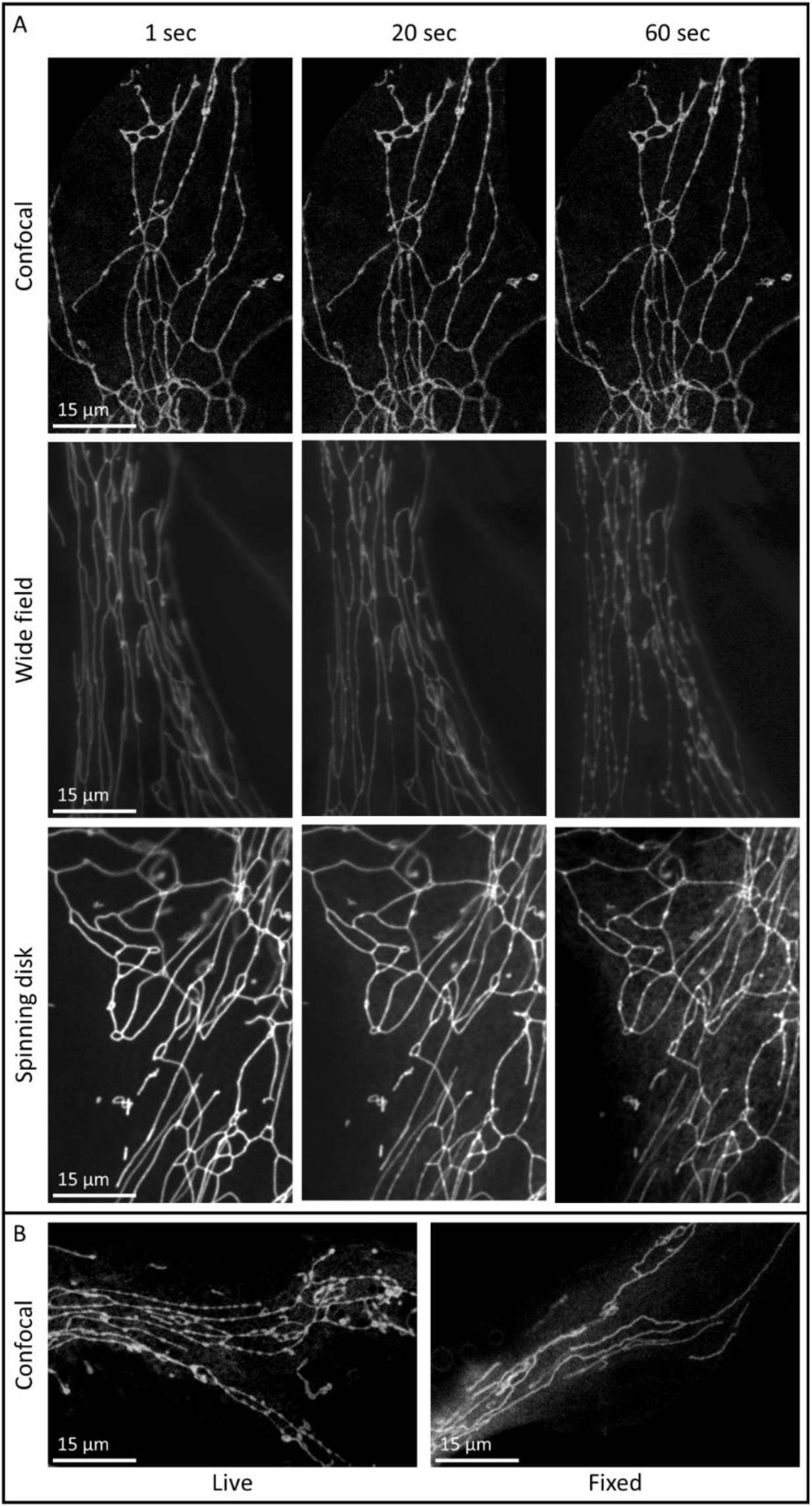
Formation of the pearl-necklace (PN) phenotype is dependent on excitation of the florescent dye by a laser.A) Scanning confocal microscopy results in immediate formation of the PN-phenotype, while the PN-phenotype formed around 20 seconds after the onset of exposure and became more severe throughout the first minute using a widefield fluorescent microscope or spinning disk confocal microscope. B) PN-phenotype is only present in living cells, not following fixation.

### 2.3 Pearl-necklace phenotype formation is dye dependent

To determine whether the PN-phenotype formation was dependent on the type of dye used for live cell imaging, cells were stained with either MitoTracker Red FM, MitoTracker CMXRos, TMRM or MitoTracker Green FM. Cells stained with the three dyes with excitation peaks in the orange-red spectrum (Red FM, CMXRos, TMRM) all presented with the PN-phenotype, while cells stained with MitoTracker Green FM did not show the PN-phenotype (Figure 4). Interestingly, this was not due to the excitation wavelength that was used. When cells stained with MitoTracker Red-FM were excited with 490 nm instead of 572 nm the cells could still form the phenotype (Figure 4). However, the observed phenotype is less severe compared to cells excited with 572, corresponding with the weaker excitation of the MitoTracker Red-FM dye at 490 nm. Conversely, cells stained with MitoTracker Green-FM that were excited with 561 nm instead of 490 nm still do not show the PN-phenotype. This indicates that the formation of the PN-phenotype is dependent on the dye used, in combination with the level of excitation.

**Figure 4:**
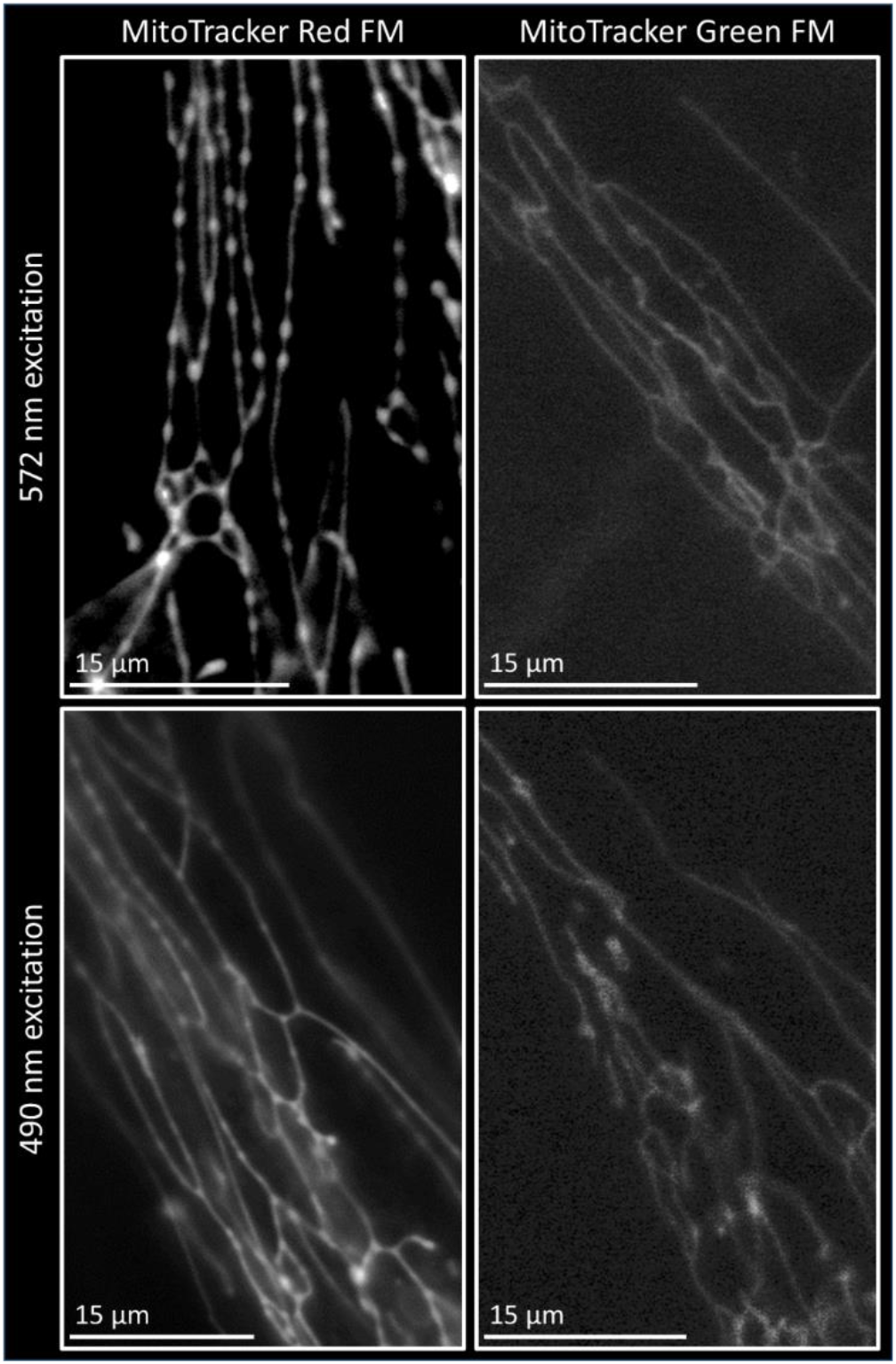
Formation of the pearl-necklace (PN) phenotype is dependent on the dye used for live staining. Cells stained with MitoTracker Red FM present with PN-structures when excited with both 572/35 nm and 490/20 nm, with a less severe PN-phenotype upon excitation with 490 nm, corresponding with the lower excitation of the MitoTracker Red FM dye at that wavelength. Cells stained with MitoTracker Green FM do not present with the PN-phenotype, independent of the used wavelength. Images made with widefield microscopy after 30 seconds of exposure.

### 2.4 Correlative serial electron tomography reveals mitochondrial structure in pearl-necklace phenotype

To assess the mitochondrial structure of the PN-phenotype in high resolution, serial electron tomography was performed. Since the PN-phenotype is reversible, the phenotype was induced in a specific area of the well and cells were immediately fixed. To confirm PN-necklace phenotype presence, fluorescent images were captured before and after induction of the phenotype (Figure 5A/B). Following the acquisition of EM images in serial sections, the induced cells could be identified based on cell shape and orientation (Figure 5A). Furthermore, mitochondria could be identified in multiple sections that overlap with regions of PN-phenotype in the fluorescent images (Figure 5C).

**Figure 5:**
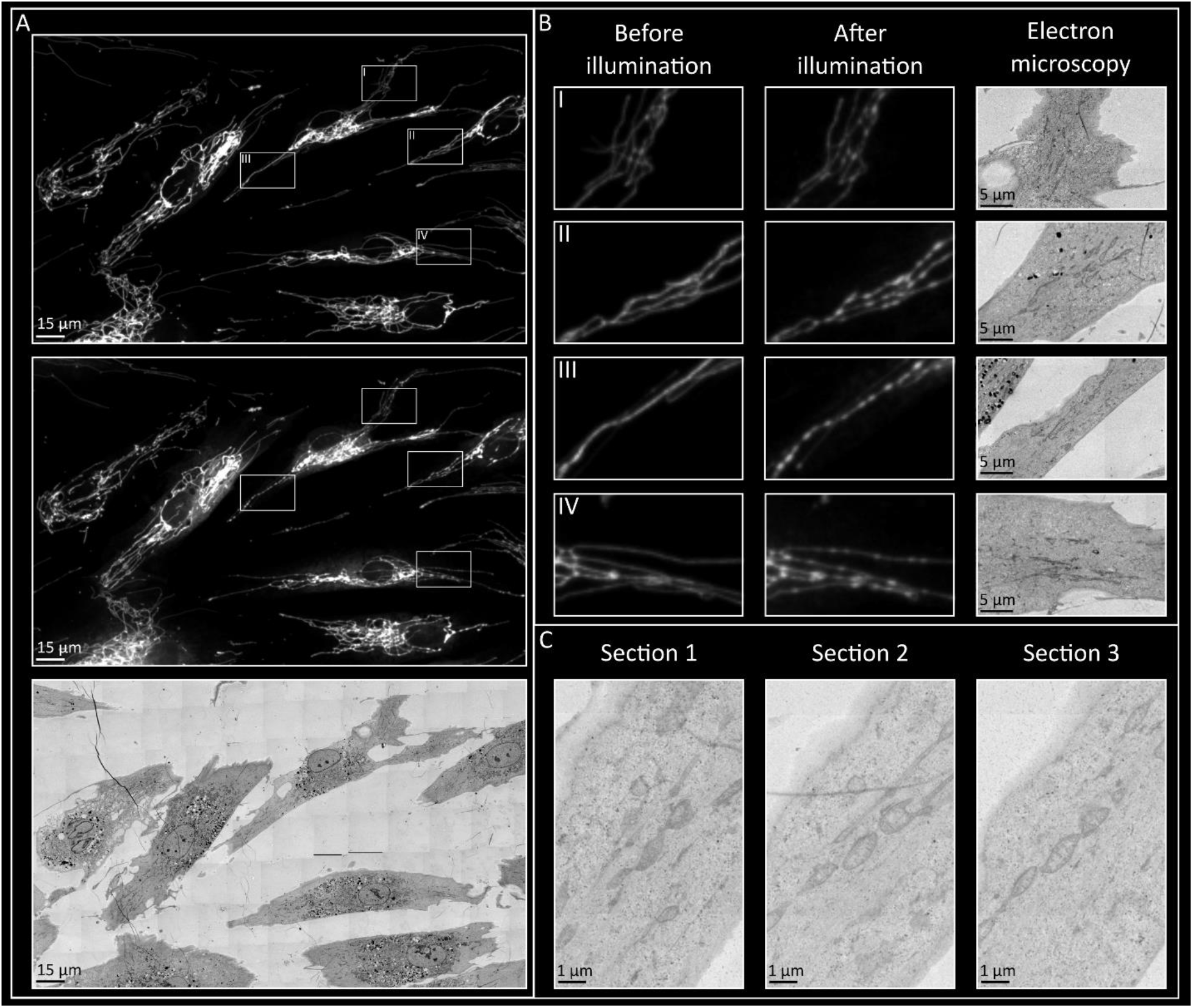
Cells with pearl-necklace phenotype, with fluorescent and electron microscopy. A) Pearl-necklace phenotype was succesfully induced with illumination and induced cell were re-identified following EM sample preparation. B) Zoomed in areas of induced cells, before and after PN-phenotype induction, with corresponding areas from the EM images. C) Mitochondria presenting with the pearl-necklace phenotype in fluorescent microscopy, could be found in multiple serial electron microscopy sections.

For serial electron tomography, the same mitochondria were localized in multiple sections and tilt series were acquired. The 3D reconstruction confirmed the presence of the PN-phenotype, characterized by alternating constricted and expanded mitochondrial regions (Figure 6). The mitochondrial double membrane remained intact in both constricted and expanded regions, indicating that no fission events had occurred at this stage (Figure 6). Similarily, cristae were observed in both the constricted and expanded areas. However, in the most expanded mitochondrial regions (Figure 6 III), cristae were present along the outer edges but absent in the middle, suggesting a somewhat altered cristae structure in the PN-phenotype. Additionally, ER and actin and/or intermediate filaments were found near the constricted areas (Figure 6), indicating mitochondrial pre-constriction by the ER, which occurs before mitochondrial fission.

**Figure 6:**
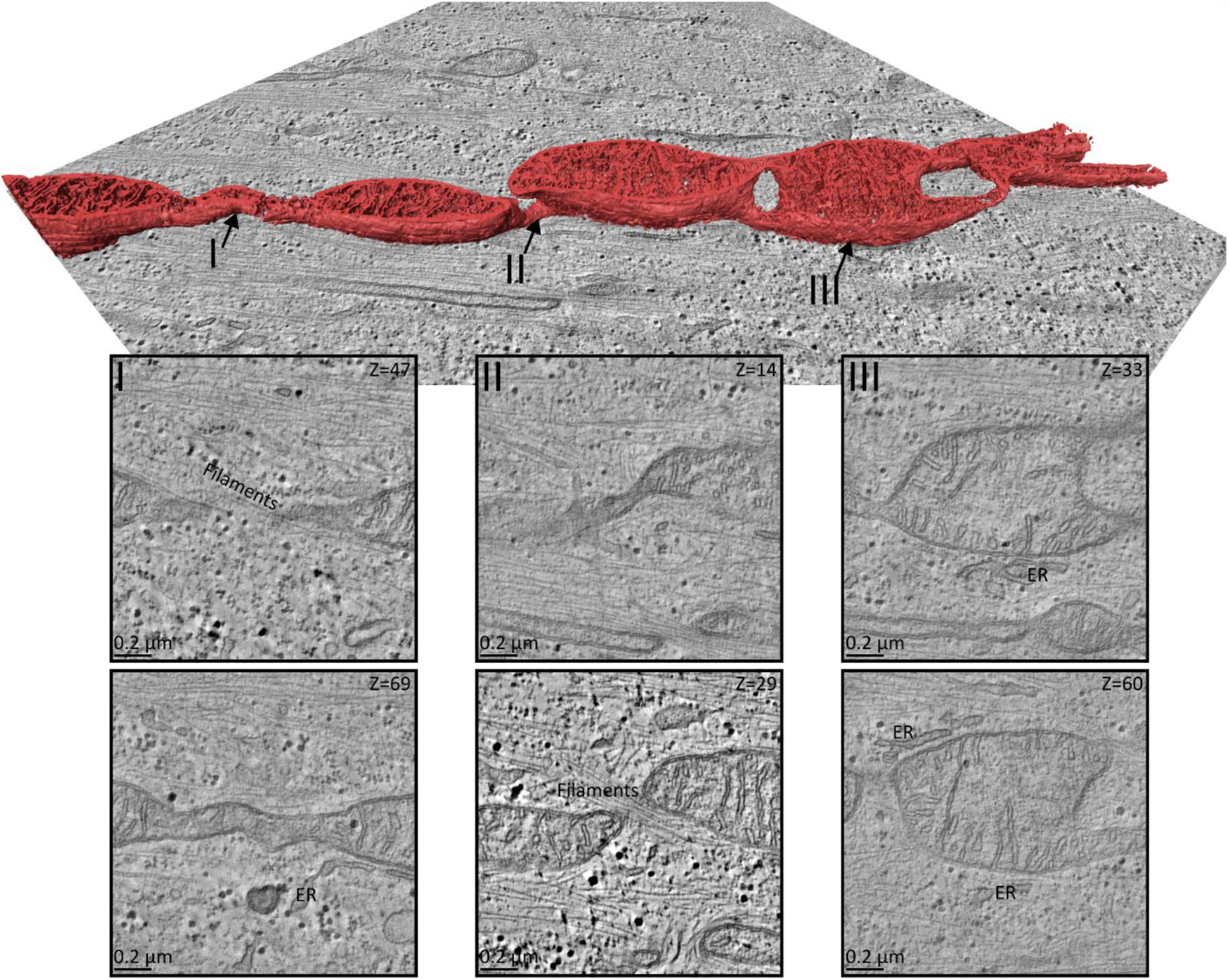
3D reconstruction of the pearl-necklace phenotype from serial electron tomography. Expanded mitochondria are alternated with constricted pieces. Cristae are present in constricted areas, although with a lower abundance than in expanded regions. Additionally, ER and filaments are present alongside constricted areas.

## 3. Discussion

Visualization of mitochondria through various live cell imaging techniques is crucial in mitochondrial dynamics research, providing insight in the structural alterations that occur due to changes in mitochondrial fission, fusion and transport. In this study, we observed a fibroblast specific pearl-necklace (PN) like phenotype, which was not observed in other cell types including HeLa cells, HUVEC cells and mesoangioblasts. This phenotype appears irrespective of the fission factor knocked down or the genetic background of the fibroblast. The pearl-necklace like phenotype was previously observed in fibroblasts obtained from patients with mutations in fission factors *DNM1L* (Drp1) and *MFF* imaged with confocal microscopy, however the origin was not investigated in these studies [20, 21]. Starvation experiments were conducted to reveal whether the PN-phenotype was the result of increased fusion or decreased fission. Mild starvation increases mitochondrial fusion through HDAC6-dependent deacetylation of mfn1 [22]. Extreme starvation on the other hand results in a hyperfused network by inhibiting phosphorylation of Drp1, which prevents recruitment to the mitochondria and thus mimicking Drp1 depletion [22]. Interestingly, mild starvation resulted in a hyperfused network but no PN-phenotype, while extreme starvation did result in the PN-phenotype. These results indicate that the PN-phenotype is caused by a decreased fission activity and not the result of increased fusion.

Strikingly, the PN-phenotype can only be observed in live cells, but not in fixed cells, and is reversible over time when light exposure is stopped, indicating that it is a dynamic adaptation related to the imaging process. Furthermore, we observed that the time it takes for the mitochondrial network to form the PN-phenotype differs between various microscopes. The PN-phenotype was most severe when using the LSCM and appeared only after 20 seconds using widefield microscopy and SDCM. LSCM generally uses a high laser intensity to visualize structures and it is a relatively slow imaging technique. The signal is received pixel for pixel by the use of a pinhole, increasing the signal-to-noise ratio and allowing imaging of a specific cell layer. However, the consequence of this technique is that the cell is exposed for a longer time to the fluorescent laser at a high intensity [23]. Contrastingly, widefield microscopy does not use a pinhole system and results in images with more background signal, especially in thicker samples. On the contrary, the imaging is significantly faster as the field of view does not need to be scanned pixel by pixel [23]. While the total laser exposure on the cells is similar to a LSCM, the exposure is shorter and less focused on a single cell layer, decreasing the exposure per mitochondrion. Spinning disk confocal microscopy (SDCM), has the advantage of an increased signal-to-noise ratio due to the use of pinholes, while simultaneously being a fast imaging method through the use of multiple spinning disks containing pinholes [23]. This results in a decrease in required exposure time compared to the LSCM. The correlation between the level of laser exposure and the severity of the PN-phenotype indicates that it is a dynamic adaptation of the mitochondrial morphology over imaging time, caused by the light exposure.

The main cause of negative cellular effect of laser exposure during live cell imaging is phototoxicity. Advanced phototoxicity is visible on a whole cell level and is characterized by changes in cell morphology, cell motility and cell death [16, 24]. While these cellular changes are generally used to determine phototoxicity, more subtle changes can be observed earlier, which also significantly affect the cell. These changes include a slowed down cell cycle and changes in cytoplasmic calcium as well as effects on mitochondrial membrane potential and dynamics [16, 17]. Considering the direct link between laser exposure and PN-phenotype severity, the onset is most likely the result of phototoxicity. The main cause of phototoxicity is the production of reactive oxygen species (ROS) as the result of a reaction between fluorophores and oxygen following laser exposure [16, 24]. Generally, there are not sufficient ROS scavengers in the cells to handle this additional ROS production [16], resulting in prolonged exposure to the produced ROS, which induces DNA and mitochondrial damage and causes cellular stress [25]. Under physiological circumstances mitochondria undergo increased amounts of fusion under light stress, to increase OXPHOS capacity and lower the load of damaged compartments [25]. Furthermore, mitochondria undergo fission to split of damaged mitochondrial parts, which can subsequently be recycled through mitophagy, to decrease the level of damage [25]. Since mitochondrial fission is substantially decreased in the Drp1 knockdown cells, this damage mitigation is largely prevented. Buildup of mitochondrial damage eventually results in apoptosis and the observed phenotype might be a cellular response initiated to prevent apoptosis. For instance, the constriction sites observed in the PN-phenotype are possibly caused by the initiation of fission at many sites simultaneously, in order to increase the chances of a successful fission event. In case the laser exposure is halted, the additional production of ROS is ceased and the present scavengers can remove the access build up, allowing the cells to recover, which is in line with the recovery of the PN-phenotype following the termination of laser exposure as observed in this study.

Phototoxicity as the result of ROS production is generally expected to be increased when exciting cells with a shorter wavelength, due to the high energy levels [26]. If this were solely the case for the PN-phenotype, it should result in an increase in the phenotype when cells are exposed to a 490 nm light source, compared to a 572 nm light source, while the opposite was observed. In fact, the formation of the PN-phenotype does not appear to be wavelength dependent. If wavelength were a determining factor, cells stained with MitoTracker Green FM, excited a 572 nm light should result in the PN-phenotype, while cells stained with MitoTracker Red FM, excited with 490 nm light should not. Instead, the PN-phenotype is dye dependent, with dyes that have excitation peaks in the orange/red spectrum presenting with the phenotype (Figure 4). Furthermore, the level of excitation of these dyes is a factor in the extent to which the PN-phenotype is formed. Red dyes excited with 490 nm results in the phenotype to a lesser extent than 572 nm, likely through the less efficient excitation of red dyes at 490 nm. Previous research has shown distinct phototoxicity in cells stained with red MitoTrackers, which was absent upon staining with various green dyes (Rhodamine123, MitoTracker Green, JC-1) [27, 28]. Additionally, the damage was less severe in presence of supplemented antioxidants, indicating that the cause of the damage was indeed ROS production [28]. This suggests that there is a chemical reaction resulting in ROS production that is specific to red dyes.

Serial electron tomography was performed to identify the effect of the PN-phenotype on the high resolution mitochondrial structure. We showed that application of correlative fluorescent and electron microscopy, in combination with serial electron tomography is a feasible technique, even for a phenotype which quickly reverses to the original state. Application of this approach resulted in the observation that the double mitochondrial membrane remained intact in the PN-phenotype. While there are reports that the inner membrane can divide separately from the outer membrane, an event that often precedes fission, this does not occur in the PN-phenotype [29, 30]. Cristae structure also appears to be altered in the most expanded PN-phenotype regions, where the cristae are not present in the mitochondrial center. This is in line with a recent study that observed stacking of cristae along the mitochondrial edges, with a hollow space in the middle, as the result of phototoxicity [18]. Additionally, the presence of ER and filaments along the constricted mitochondrial regions indicate that the constrictions are the result of ER-mediated mitochondrial pre-constriction, which normally happens before mitochondrial fission [3, 29]. This would be in line with the previously mentioned hypothesis that fission is initiated at many sites simultaneously in these cells, to mitigate the damaging effect of the induced phototoxicity. However, due to the Drp1 depletion, fission cannot properly take place resulting in the observed PN-structures.

The PN-phenotype was found to be a change in mitochondrial morphology dependent on the amount of laser exposure and the choice of dye. This shows that while live cell imaging is crucial to study dynamic cellular structures, the obtained results are not always representative of the system under physiological circumstances. Especially in dynamic organelles, which are capable of quickly adapting to presented stimuli, the possible effects of phototoxicity should be taken under consideration when analyzing obtained results, to prevent misinterpretation. Additionally, these results show that the choice of microscope and live cell staining should be carefully considered when designing live cell imaging experiments.

## 4. Methods

### 4.1 Cell culture

Normal human dermal fibroblasts (nHDFs), c0388, c0407, c2244, HeLa cells and HepG2 cells were cultured in DMEM medium (Gibco, 41966029) supplemented with 10% fetal bovine serum (FBS, Takara, 631106) and 0.1% PenStrep and maintained at 37 °C in a humidified incubator with 5% CO_2_. For HUVEC cells EGM-2 endothelial cell growth medium (Lonza, CC-3162) was used and cells were kept under the same conditions as mentioned before. Mesoangioblasts (mabs) were cultured in IMDM (Gibco, 12440053) supplemented with sodium-pyruvate, glutamine, non-essential amino acids, insulin transferase selenium X, 0.2% 2-mercaptoethanol, 10% FBS, 5 ng/mL FGF2 and 0.1% gentamycin. These cells were incubated at 37 °C in a humidified incubator with 5% CO_2_ and 4% O_2_. Cells were split at 70% confluency and were seeded 24 hours before transfection onto 8 well µ-Slides (Ibidi, 80826), with a density of 1000 (fibroblasts & mabs) or 5000 cells (HeLa, HepG2 and HUVEC) per well.

### 4.2 Transient knockdown

An esiRNA-mediated knockdown was created for *DNM1L* (Drp1), *MFF, MIEF1, MIEF2, FIS1* and a non-targeting control (GFP). T7 promotor sequence containing primers were used to amplify around 600bp of the respective transcripts with standard PCR amplification using BIOTAQ DNA Polymerase (Meridian Bioscience, BIO-21040) and cDNA as a template. cDNA was made with the qScript cDNA Supermix kit (Quantabio, 95048-500), with 400 ng RNA as input. dsRNA was synthesized from the PCR products using the MEGAscript Kit (Ambion, AM1334) according to manufacturer protocol. For esiRNA production, dsRNA was incubated with Shortcut RNase III (NEB, M0245S) according to manufacturer instructions, with 0.75 units of enzyme per 1 µg dsRNA. To induce knockdown, cells were transfected with esiRNAs. In short, esiRNAs were combined with RNAiMAX (Invitrogen, 13778075) in OptiMEM medium (Gibco, 11058021) with a concentration of 50 ng esiRNA/ml culture medium. Further experiments were performed 72 hours after transfection.

### 4.3 Knockdown efficiency

Normal Human Dermal Fibroblasts were seeded in a 6-well plate with a density of 10,000 cell per well and transfected with esiRNAs. Following 72h of incubation, RNA was isolated with the RNeasy Mini Kit (Qiagen, 74104) and the RNAse-Free DNase Set (Qiagen, 79254) according to manufacturer instructions. cDNA was synthesized using the qScript cDNA Supermix kit (Quantabio, 95048-500), with 400 ng RNA as input. qPCR was performed using 1x SensiMixTM SYBR® (Bioline Meridian, QT615-05 2x stock) and a final primer concentration of 0.5 μM, on the LightCycler® 480 II (Roche, 0501524300). The knockdown efficiency was calculated using the ΔΔCt between knockdown and control samples, using TBP for normalization. Significance of the relative differences was determined using an independent-sample T-test (R version 4.1.3).

### 4.4 Live staining and fixation

Following 72h of knockdown mitochondria were incubated with the respective culture medium containing MitoTracker Red FM (300 nM, Invitrogen, M22425), MitoTracker Red CMXRos (50 nM, Invitrogen, M7512), MitoTracker Green FM (300 nM, Invitrogen, M7514) or TMRM (100 nM, Invitrogen, T668) for 30 minutes at 37 °C. Cells were washed three times in culture medium and were placed in culture medium without phenol red (Gibco, 21063029), to decrease background signal during imaging. For experiments performed on fixed cells, the cells were stained with 50 nM MitoTracker Red CMXRos for 30 minutes at 37 °C, followed by washing with culture medium and fixation using 3.7% paraformaldehyde for 30 minutes. After fixation the cells were washed in PBS before imaging.

### 4.5 Microscopy

Unless mentioned otherwise, imaging was performed with a Leica TCS SPE confocal system (DMI4000B microscope, Leica LAS-AF software) with a PMT detector and a 63x oil objective (NA 1.3). Excitation lasers with wavelengths of 488 and 561 nm were used to visualize the mitochondria. For time-lapsed images, an image of the same plane was made every three seconds and cells were continuously exposed to the respective laser. For the microscope comparison, a FEI CorrSight, equipped with both a widefield and a spinning-disc module was used. Here, a 40x air objective (NA 0.9) and excitation bandwidth of 490/20 nm or 572/35 nm and an emission DAPI/FITC/TxRed Tripleband HC Filter Set were used for the widefield module. For the spinning disk module excitation lasers with wavelengths of 488 and 561 nm were used. FEI MAPS software, in combination with Live Acquisition (LA) software was used to control the microscope and capture time-lapsed images, with an exposure time of 50 ms (widefield module) or 200 ms (spinning disk module).

For the phenotype quantification, between 70 and 200 cells were randomly selected and assessed using a Leica DMI4000B microscope. For each cell, the presence/absence of the pearl-phenotype was determined. When the pearl-necklace phenotype was only present in one to two mitochondrial extremities, the cell was classified as containing an ‘intermediate’ phenotype. On the other hand, when the pearl-necklace phenotype was present in more than two extremities within one cell, this cell was classified as ‘yes’ for containing the pearl-necklace phenotype and a complete absence was classified as ‘no’. Differences between gene knockdowns were determined with an independent sample T-test (R version 4.1.3).

### 4.6 Electron tomography

For electron tomography, the pearl-necklace phenotype was induced in nHDFs cultured on a µ-Slide with a grid (Ibidi, 80826-G500). This was done with a FEI CorrSight, using the widefield module and a 20x objective (NA 0.8). Cells were exposed to a 561 nm laser for one minute and presence of the pearl-necklace phenotype was confirmed. To prevent reversal of the phenotype, cells were fixed with 2.5% Glutaraldehyde in 0.1M phosphate buffer for 24 h at 4 °C. Then cells were washed with 0.1 M cacodylate buffer and postfixed with 1% osmium tetroxide in the same buffer containing 1.5% potassium ferricyanide for 1 h in the dark at 4 °C. Samples were dehydrated in ethanol, infiltrated with Epon resin for 2 days, embedded in the same resin and polymerized at 60 ºC for 48 h. Ultrathin serial sections with a thickness of 150 nm were cut using a Leica Ultracut UCT ultramicrotome (Leica Microsystems Vienna) and mounted on Formvar-coated copper slot grids. Before staining with 2% uranyl acetate in water and lead citrate, 10nm BSA-gold fiducials were adsorbed to the grid.

Overview images of all sections were made with a FEI Tecnai G2 Spirit BioTWIN electron microscope at 2900x magnification using the EMmesh software [31]. Mitochondria that presented the PN-phenotype, both on EM and corresponding fluorescent images, and were present in multiple of the serial sections were selected for tomography. For selected positions 4800x magnified images were made, before a single-axis tilt series were recorded with a FEI Tecnai G2 Spirit BioTWIN. For each tilt series, 121 images were recorded between -65 and 65 degrees, with 1 degree increments. Images were recorded with an Eagle 4kx4k CCD camera at binning 2 (Thermo Fisher Scientific, The Netherlands), a 6800x magnification resulting in a pixelsize of 3.25 nm. Tilt series were reconstructed and tomograms form serial sections were aligned with IMOD and 3D reconstructions of PN-phenotype mitochondria were segmented and visualized with AMIRA.

## Competing interests

No competing interests declared

## Funding

This research received no specific grant from any funding agency in the public, commercial or not-for-profit sectors.

## Notes

### Competing Interest Statement

The authors have declared no competing interest.

