## Supplemental information for "The curse of the red pearl: a fibroblast specific pearl-necklace mitochondrial phenotype caused by phototoxicity"

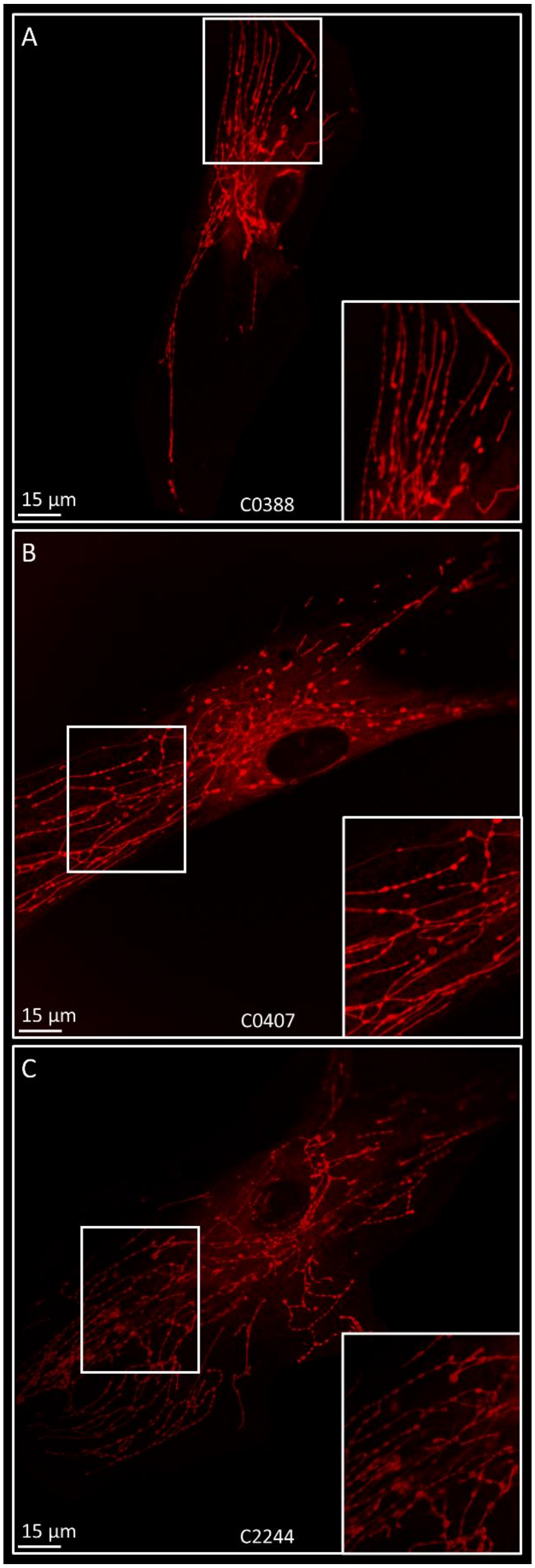


*Figure S1: The pearl-necklace phenotype is present in multiple dermal fibroblast cell lines. A) c0388 cells. B) c0407 cells. C) c2244 cells.*

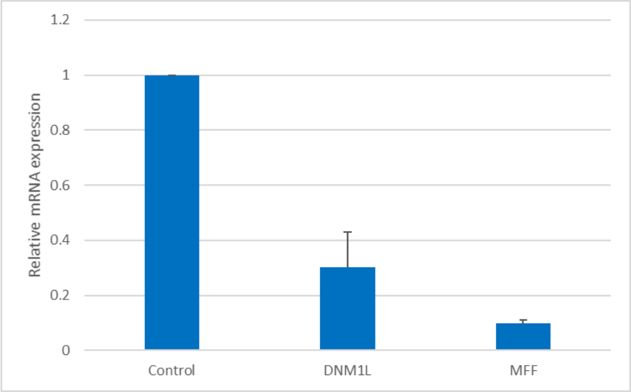


*Figure S2: Knockdown efficiency of DNM1L (Drp1) and MFF knockdown in normal human dermal fibroblasts. Both knockdown conditions show at a decrease in mRNA expression.*

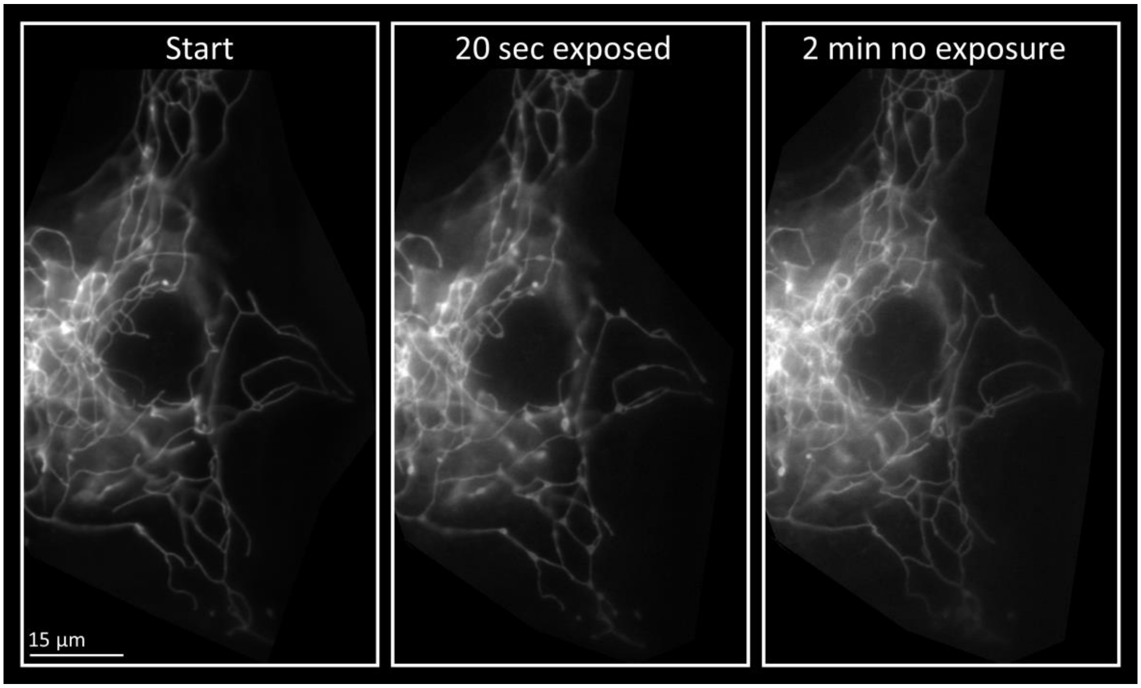


*Figure S3: Following formation of the pearl-phenotype, the mitochondrial morphology can be reversed by discontinuation of the laser excitation.*
